# RNF25 is activated as a response to amino acid starvation-induced ribosome collisions in competition with GCN2

**DOI:** 10.64898/2026.04.27.721104

**Authors:** Ivan Kisly, Ivo Zemp, Ulrike Kutay

## Abstract

Surveillance of mRNA translation relies on a suite of ribosome-associated quality control pathways. Recently, a novel pathway induced by trapping of translation factors in the ribosomal A-site has been described, involving ubiquitination of RPS27A/eS31 by the human E3 ubiquitin ligase RNF25. Here, we show that not only ribosome-stalling by low doses of translation inhibitors, but also amino acid starvation induces RPS27A/eS31 ubiquitination, identifying a natural trigger of RNF25 activation. Even under optimal growth conditions, RNF25 senses and resolves transient ribosome stalls. RPS27A/eS31 ubiquitination specifically depends on the ribosome collision sensor GCN1, a known cofactor of GCN2 involved in the integrated stress response. RNF25 and GCN2 both possess a GCN1-binding RWD domain, indicating a competitive relationship, with GCN2 acting as a negative regulator of RNF25 activation. Although both RNF25 and GCN2 respond to amino acid starvation, RPS27A/eS31 ubiquitination by RNF25 is not required for GCN2 activation, showing that both act in independent pathways. We propose that the RNF25 pathway acts as a first line of defence to resolve ribosome collisions, outcompeted by GCN2 binding to GCN1 under acute stress.

## INTRODUCTION

Protein synthesis is a fundamental biological process in which ribosomes translate genetic information encoded in mRNA into proteins. Since proteins participate in every aspect of cell life, alterations in their synthesis can disturb cellular homeostasis. For example, defects in the resolution of stalled or collided ribosomes have been associated with a wide range of pathologies, such as neurodegeneration and mitochondrial dysfunction (Choe *et al*, 2016; Chu *et al*, 2009; Defenouillere *et al*, 2016; Geng *et al*, 2024; Izawa *et al*, 2017; Martin *et al*, 2020; Udagawa *et al*, 2021; Wu *et al*, 2019b). To avoid such negative scenarios, cells have evolved a vast number of surveillance pathways that control protein synthesis (Chang & Choe, 2026; Ford *et al*, 2024; Joazeiro, 2019; Park *et al*, 2021; Yip & Shao, 2021).

Ribosome stalling can be caused by a variety of conditions, such as translation inhibitors, mRNA damaging reagents or UV radiation, the production of aberrant mRNAs, the presence of specific mRNA stalling motifs, or amino acid starvation (Chang & Choe, 2026; Park *et al*., 2021). Three major pathways recognize and handle ribosome collisions: the ribosome-associated quality control (RQC), the integrated stress response (ISR) and the ribotoxic stress response (RSR) (Chang & Choe, 2026; Park *et al*., 2021). The RQC pathway serves as a first line of defence dealing with a basal level of collisions that may occur even under optimal growth conditions. Here, collided ribosomes are recognized by the core RQC factor ZNF598 (or the yeast homolog Hel2) (Juszkiewicz *et al*, 2018; Juszkiewicz & Hegde, 2017; Sundaramoorthy *et al*, 2017). Independently of that, the collision sensor EDF1 recruits GIGYF2-4EHP to the stalled ribosomes, which blocks translation initiation to reduce usage of the affected mRNAs (Juszkiewicz *et al*, 2020a; Sinha *et al*, 2020). These events promote the engagement of factors responsible for the splitting of ribosomes into 40S and 60S subunits as well as degradation of aberrant mRNAs and nascent polypeptides. Upon more global translational stress causing more persistent collisions, the ISR pathway is additionally activated through the recognition of collided ribosomes by GCN1 and subsequent recruitment of the kinase GCN2 to the stalling ribosome. Active GCN2 phosphorylates eIF2α, causing both the inhibition of translation and transcriptional activation of stress response genes (Castilho *et al*, 2014; Darnell *et al*, 2018; Harding *et al*, 2019; Hinnebusch, 2005; Ishimura *et al*, 2016; Marton *et al*, 1993; Pochopien *et al*, 2021; Wu *et al*, 2020; Zhou *et al*, 2025). Finally, upon severe and persistent translational stress, the RSR pathway is induced by the MAP3K ZAKα, which becomes activated on collided ribosomes and initiates the activation of pro-inflammatory and pro-apoptotic pathways (Huso *et al*, 2026; Sinha *et al*, 2024; Stoneley *et al*, 2022; Vind *et al*, 2020). It is important to note, that although these three pathways may be considered independent processes, they have complex antagonistic as well as cooperative relations with each other (Chang & Choe, 2026; Kim *et al*, 2024; Park *et al*., 2021; Pochopien *et al*., 2021).

An important role in surveillance pathways is assigned to protein ubiquitination, which facilitates protein degradation but also has many regulatory functions (Dougherty *et al*, 2020; Ford *et al*., 2024). For example, recent studies showed that regulatory ribosomal ubiquitination is a crucial step in the activation of the RQC pathway (Juszkiewicz & Hegde, 2017; Sundaramoorthy *et al*., 2017). Collided ribosomes induce ubiquitination of the ribosomal proteins RPS10/eS10 and RPS20/uS10 by the E3 ubiquitin ligase ZNF598, which is antagonized by deubiquitinases (DUBs) USP21 and OTUD3 (Garshott *et al*, 2020; Ikeuchi *et al*, 2019; Juszkiewicz *et al*., 2018; Juszkiewicz & Hegde, 2017; Sundaramoorthy *et al*., 2017). Ubiquitination of ribosomal proteins in RQC is needed to recruit the ASCC complex, which facilitates splitting of stalled ribosomes (Best *et al*, 2023; Hashimoto *et al*, 2020; Juszkiewicz *et al*, 2020b; Matsuo *et al*, 2017; Matsuo *et al*, 2023; Narita *et al*, 2022). However, despite advances in understanding how ubiquitination acts in translational stress, many aspects and targets of such modifications remain uncharacterized, and regulatory ribosomal ubiquitination remains an important topic for both basic science and medical research.

Recently, we discovered a novel translation-dependent mono-ubiquitination event targeting RPS27A/eS31 in human cells (Fig 1A) (Montellese *et al*, 2020). We demonstrated that RPS27A/eS31 is modified on lysine K113, which is counteracted by the DUB USP16 (Montellese *et al*., 2020). Further work in the field then revealed that RPS27A/eS31 ubiquitination is mediated by the E3 ligase RNF25 as a part of a novel RQC pathway, which is induced in response to treatment of cells with the translation inhibitors ternatin-4, SRI-41315 or NVS1.1 (Gurzeler *et al*, 2023; Oltion *et al*, 2023). These drugs bind the translation factors eEF1A (in case of ternatin-4) and eRF1 (in case of SRI-41315 and NVS1.1), respectively, and trap them in the A-site of the ribosome (Gurzeler *et al*., 2023; Oltion *et al*., 2023). Affected ribosomes stall on mRNA, which causes ribosome collisions that are sensed by GCN1. This leads to RPS27A/eS31 ubiquitination and subsequent recruitment of the E3 ubiquitin ligase RNF14 for ubiquitination and clearance of the trapped translation factors from the ribosome (Gurzeler *et al*., 2023; Oltion *et al*., 2023).

**Figure 1.**
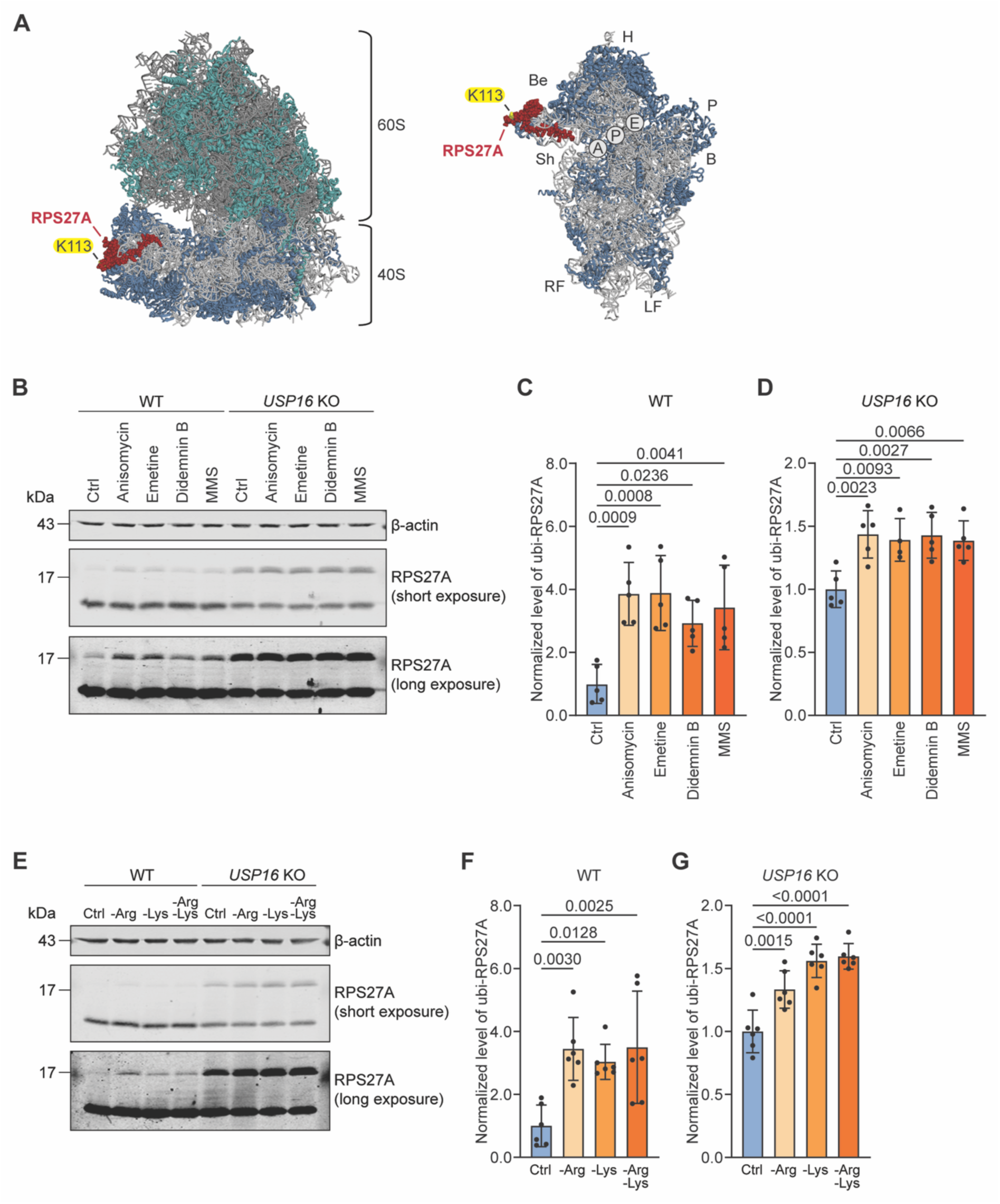
Ubiquitination of RPS27A/eS31 is induced by conditions that cause ribosome stalling including amino acid starvation. **(A)** Structural models of the human 80S ribosome (left) and of the respective 40S subunit (right) (PDB ID: 6QZP, (Natchiar *et al*, 2017)). Ribosomal protein RPS27A/eS31 (red) as well as its K113 (yellow) are indicated. 80S is shown from the P-stalk side, 60S subunit (rRNA in dark grey and proteins in teal) and 40S subunit (rRNA in light grey and proteins in blue) are indicated. On the right, 40S is shown from the subunit interface side. Positions of A-, P-, and E-sites as well as major structural elements of the 40S are indicated (B, body; Be, beak; H, head; LF, left foot; RF, right foot; Sh, shoulder). **(B)** Induction of RPS27A/eS31 ubiquitination by diverse ribosome-stalling compounds. HeLa WT and *USP16* KO cells were cultivated in the absence or presence of the following drugs: 0.19 µM anisomycin (30 min), 1.8 µM emetine (30 min), 0.5 µM didemnin B (30 min, 0.25 µg/ml MMS (60 min). Cell extracts were analysed by immunoblotting with the indicated antibodies. **(C, D)** Quantification of levels of ubiquitinated RPS27A/eS31 in HeLa WT C) and *USP16* KO (D) lysates from (B), expressed as the ratio between ubi-RPS27A/eS31 and total RPS27A/eS31 (ubi-RPS27A/eS31 + unmodified RPS27A/eS31) and normalized to samples from untreated cells (N ≥ 4, mean ± SD, one-way ANOVA and post hoc Dunnett’s test; p-values are indicated). **(E)** Amino acid starvation enhances RPS27A/eS31 ubiquitination. HeLa WT and *USP16* KO cells were grown for 6 h in control conditions (Ctrl, no starvation), or in the absence of arginine (-Arg), lysine (-Lys) or both (-Arg-Lys). Cell extracts were analysed by immunoblotting with the indicated antibodies. **(F, G)** Quantification of levels of ubiquitinated RPS27A/eS31 in HeLa WT (F) and *USP16* KO (G) lysates from (E), expressed as the ratio between ubi-RPS27A/eS31 and total RPS27A/eS31 (ubi-RPS27A/eS31 + unmodified RPS27A/eS31) and normalized to samples from non-starved cells (N = 6, mean ± SD, one-way ANOVA and post hoc Dunnett’s test, p-values are indicated.

Here, we have investigated the triggers and the underlying mechanism of the RNF25 pathway in more detail. Our analysis revealed that sub-inhibitory concentrations of several translation inhibitors induce activation of RNF25. We identify amino acid starvation as a physiological stalling condition that induces ubiquitination of RPS27A/eS31. Both compound- and starvation-induced RPS27A/eS31 ubiquitination is specifically dependent on the collision sensor GCN1, but not on EDF1 or ZAKα, establishing RPS27A/eS31 ubiquitination as part of a distinct pathway. Since GCN1 is known to be involved in the ISR, we evaluated the relation of RPS27A/eS31 ubiquitination in relation to stress signalling by GCN2. Exploiting cells harbouring a ubiquitination-deficient K113R mutant, we show that RNF25 is not required for GCN2 activation, establishing them as factors acting independently downstream of GCN1. In support of this model, RNF25 gets hyperactivated in the absence of GCN2, suggesting that both pathways co-exist as two parallel pathways dependent on GCN1, guarding cells from ribosome stalling.

## RESULTS

### Ubiquitination of RPS27A/eS31 is induced by conditions that cause ribosome stalling

Several recent studies have described ubiquitination of RPS27A/eS31 as a part of a protein synthesis quality control pathway activated in the presence of drugs that trap eEF1A or eRF1 in the A-site of the ribosome, causing stalling and subsequent collision of ribosomes (Gurzeler *et al*., 2023; Oltion *et al*., 2023). Here, we aimed at better understanding the mechanism of RPS27A/eS31 ubiquitination. First, we tested whether we could detect ubiquitination of RPS27A/eS31 upon treatment of HeLa WT cells with ternatin, a commercially available but slightly less potent analogue of ternatin-4 (Carelli *et al*, 2015). Quantitative immunoblotting indeed revealed increased levels of ubiquitinated RPS27A/eS31 after ternatin treatment (Fig S1A, S1B), consistent with previous work (Oltion *et al*., 2023). However, the levels of ubiquitinated RPS27A/eS31 in WT cells are relatively low, hampering a robust quantitative assessment of this modification due to a low signal-to-noise ratio (Fig S1A, S1B). To confirm our results and to circumvent this issue, we therefore analysed RPS27A/eS31 ubiquitination in HeLa *USP16* KO cells. Due to the lack of USP16, the DUB for RPS27A/eS31 (Montellese *et al*., 2020), these cells naturally contain higher levels of ubiquitinated RPS27A/eS31, which are further increased upon induction of stalling, facilitating quantitative analyses (Fig S1A, S1C). Indeed, ternatin increased the levels of ubiquitinated RPS27A/eS31 in these cells about 1.5-fold.

To understand whether other ribosome stalling conditions can also induce ubiquitination of the RPS27A/eS31, HeLa WT and *USP16* KO cells were treated with drugs known to stall ribosomes through various mechanisms, i.e. anisomycin, emetine, didemnin B and methyl methanesulfonate (MMS) (Fig 1B-D). Anisomycin binds to the 60S ribosomal subunit and prevents peptide bond formation (Garreau de Loubresse *et al*, 2014). Emetine binds to the 40S ribosomal subunit and blocks tRNA translocation (Wong *et al*, 2014). Didemnin B acts similarly to ternatin, trapping the translation elongation factor eEF1A on the ribosome (Juette *et al*, 2022). MMS alkylates mRNA, which creates bulky mRNA adducts that cause ribosome stalling by interfering with aminoacyl-tRNA accommodation (Thomas *et al*, 2020; Yan & Zaher, 2021). All these conditions induced ubiquitination of RPS27A/eS31 in both WT and *USP16* KO cells (Fig 1B-D). This indicates that ubiquitination of RPS27A/eS31 is a general response to drug-induced ribosome stalling, irrespective of the mechanism by which stalling is achieved.

Next, we wished to explore which physiological conditions can induce RPS27A/eS31 ubiquitination. It is known that amino acid starvation can cause stalling of ribosomes by reducing the levels of the respective aminoacyl-tRNAs (Darnell *et al*., 2018; Worpenberg *et al*, 2025; Zhou *et al*., 2025). We therefore cultivated WT and *USP16* KO cells in the absence of arginine, lysine or both. Immunoblot analysis revealed that in both cell lines, levels of ubiquitinated RPS27A/eS31 increased in the absence of amino acids (Fig 1E-G).

Altogether, these results indicate that ubiquitination of RPS27A/eS31 is induced by a variety of conditions that cause ribosome stalling. Importantly, also physiological shortage of amino acids as occurring during amino acid starvation can trigger RPS27A/eS31 ubiquitination.

### Ubiquitination of RPS27A/eS31 depends on RNF25 and GCN1

Recent studies have identified RNF25 as a candidate E3 ubiquitin ligase responsible for the ubiquitination of the RPS27A/eS31 (Gurzeler *et al*., 2023; Oltion *et al*., 2023). To investigate whether the elevated levels of RPS27A/eS31 ubiquitination in *USP16* KO cells induced by translation inhibitors or amino acid starvation can indeed be attributed to the action of RNF25, we generated *USP16/RNF25* double knockout (DKO) cell lines from parental HeLa *USP16* KO cells using CRISPR/Cas9. The *RNF25* deletion was confirmed by both genotyping and immunoblotting (Fig S2A, S2B). In addition, we validated inactivation of the RNF25 pathway by treating cells with SRI-41315, which traps eRF1 on ribosomes and leads to its RNF25-dependent ubiquitination and subsequent degradation (Oltion *et al*., 2023). As expected, knockout of *RNF25* prevented the degradation of eRF1 (Fig S2B), confirming that our knockout was functional. Compared to the parental *USP16* KO cells, *USP16/RNF25* DKO cells failed to induce ubiquitination of RPS27A/eS31 upon amino acid starvation (Fig 2A, 2B). Thus, the rise in the levels of ubiquitinated RPS27A/eS31 upon ribosome stalling in response to amino acid starvation depends on RNF25.

**Figure 2.**
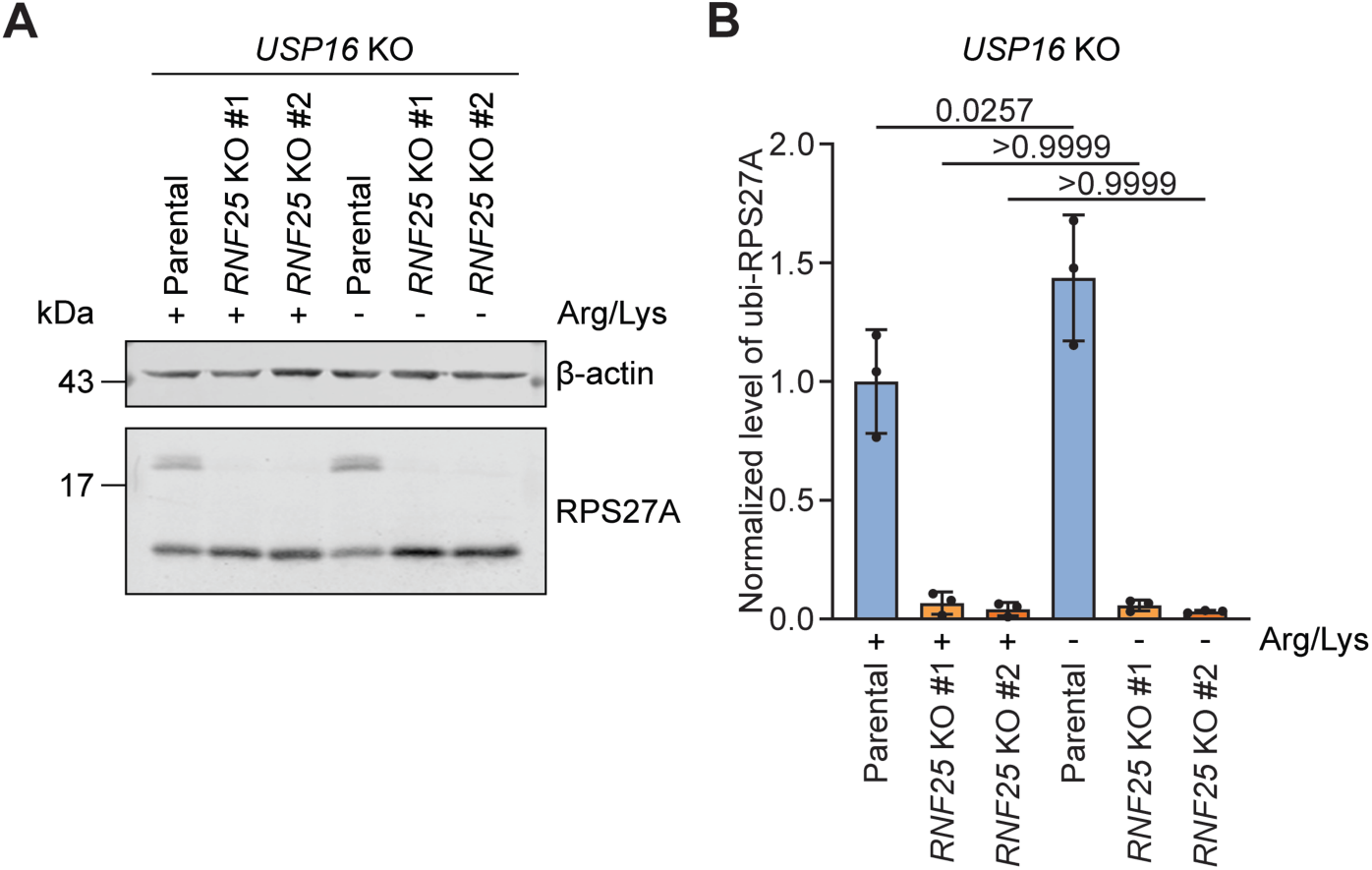
Ubiquitination of RPS27A/eS31 depends on RNF25. **(A)** Analysis of RPS27A/eS31 ubiquitination in HeLa *USP16* KO cells upon deletion of *RNF25*. The *RNF25* gene was deleted from *USP16* KO cells by CRISPR/Cas9. The parental *USP16* KO cells and two clones of *USP16*/*RNF25* double knockout (DKO) cells were cultivated for 6 h in the presence (+) or absence (-) of arginine and lysine. Cell extracts were analysed by immunoblotting with the indicated antibodies. **(B)** Quantification of levels of ubiquitinated RPS27A/eS31 in cell lysates from (A), expressed as the ratio between ubi-RPS27A/eS31 and total RPS27A/eS31 (ubi-RPS27A/eS31 + unmodified RPS27A/eS31) and normalized to samples from non-starved parental *USP16* KO cells (N = 3, mean ± SD, one-way ANOVA and post hoc Tukey’s test, p-values are indicated).

Several proteins have been described as sensors of stalled/collided ribosomes: ZNF598, GCN1, EDF1 and ZAKα (Huso *et al*., 2026; Juszkiewicz *et al*., 2018; Pochopien *et al*., 2021; Sinha *et al*., 2020) (Fig 3A). We have previously demonstrated that ZNF598 is not involved in RPS27A/eS31 ubiquitination (Montellese *et al*., 2020), although it mediates the ubiquitination of other ribosomal proteins in response to ribosome stalling (Juszkiewicz & Hegde, 2017; Sundaramoorthy *et al*., 2017). In contrast, the RNF25 pathway is activated through the recognition of collided ribosomes by GCN1 (Gurzeler *et al*., 2023; Oltion *et al*., 2023). To define whether only GCN1, or also EDF1 and ZAKα, are involved in ubiquitination of RPS27A, we depleted these factors in *USP16* KO cells using siPOOLs (Fig 3B-D), followed by ternatin treatment to induce ribosome stalling and to activate the RNF25 pathway. The knockdown of GCN1 led to reduced levels of ubiquitinated RPS27A/eS31 under control conditions, and no increase in ubiquitination upon ribosome stalling, as expected. In contrast, depletion of EDF1 and ZAKα had no effect on the levels of ubiquitinated RPS27A/eS31 compared to control conditions, in the presence or absence of ternatin (Fig 3B-D). These data indicate that GCN1 is required for RNF25-dependent RPS27A/eS31 ubiquitination, whereas EDF1 and ZAKα are not.

**Figure 3.**
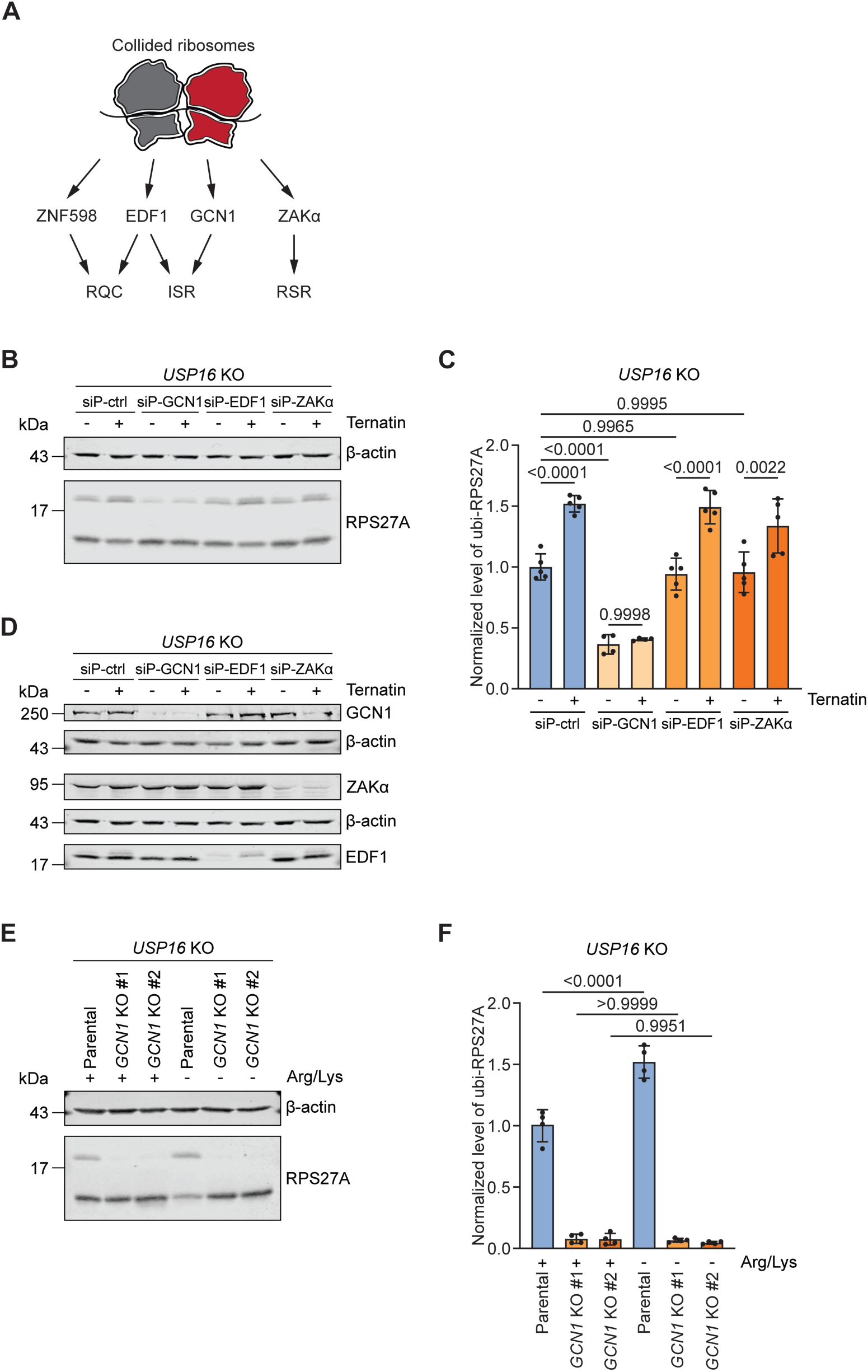
Ubiquitination of RPS27A/eS31 depends on GCN1 but not on EDF1 or ZAKα. **(A)** Schematic depiction of various sensors that recognize collided ribosomes in three major quality control pathways: ribosome-associated quality control (RQC), integrated stress response (ISR) and ribotoxic stress response (RSR). **(B)** Analysis of RPS27A/eS31 ubiquitination in HeLa *USP16* KO cells upon depletion of GCN1, EDF1 or ZAKα. *USP16* KO cells were treated for two times 48 h with siPOOLs (control or targeting the mRNAs of GCN1, EDF1 or ZAKα). Prior to harvesting, cells were cultivated for 4 h in the absence (-) or presence (+) of 50 nM ternatin. Cell extracts were analysed by immunoblotting with the indicated antibodies. **(C)** Quantification of levels of ubiquitinated RPS27A/eS31 in cell lysates from (B), expressed as the ratio between ubi-RPS27A/eS31 and total RPS27A/eS31 (ubi-RPS27A/eS31 + unmodified RPS27A/eS31) and normalized to samples from untreated control cells (N ≥ 4, mean ± SD, one-way ANOVA and post hoc Tukey’s test, p-values are indicated). **(D)** Validation of knockdown experiments from (B). The efficiency of downregulation was analysed by immunoblotting of cell extracts with the indicated antibodies. **(E)** Analysis of RPS27A/eS31 ubiquitination in HeLa *USP16* KO cells upon deletion of *GCN1*. The *GCN1* gene was deleted from *USP16* KO cells by CRISPR/Cas9. The parental *USP16* KO cells and two clones of *USP16*/*GCN1* DKO cells were cultivated for 6 h in the presence (+) or absence (-) of arginine and lysine. Cell extracts were analysed by immunoblotting with the indicated antibodies. **(F)** Quantification of levels of ubiquitinated RPS27A/eS31 in cell lysates from (E), expressed as the ratio between ubi-RPS27A/eS31 and total RPS27A/eS31 (ubi-RPS27A/eS31 + unmodified RPS27A/eS31) and normalized to samples from non-starved parental *USP16* KO cells (N = 4, mean ± SD, one-way ANOVA and post hoc Tukey’s test, p-values are indicated).

To test whether GCN1 is also involved in starvation-induced RPS27A/eS31 ubiquitination, we generated HeLa *USP16/GCN1* DKO cell lines (Fig S2C, S2D). As expected, compared to the parental *USP16* KO cells, RPS27A/eS31 modification was impaired in the *USP16/GCN1* DKO. Importantly, amino acid starvation failed to induce RPS27A/eS31 modification in the absence of GCN1 (Fig 3E, 3F). This confirms that GCN1 is required for RPS27A/eS31 ubiquitination in response to decreased availability of amino acids.

Taken together, our results indicate that the ubiquitination of RPS27A/eS31 depends on the recognition of collided ribosomes by GCN1 and recruitment of RNF25. Other collision sensors, such as EDF1 and ZAKα, seem dispensable for RPS27A/eS31 ubiquitination.

### Ubiquitination of RPS27A/eS31 is needed to support translation

To understand how inactivation of the RNF25 pathway influences translation, we analysed polysome profiles of HeLa WT cells upon knockdown of RNF25 (Fig 4A-C). Interestingly, in the absence of RNF25, cells displayed slightly decreased levels of 80S monosomes, but the amount of polysomes was similar to control cells, resulting in an increased polysome/monosome (P/M) ratio. This might indicate that RNF25 is needed to support clearance of transiently stalled ribosomes already under optimal growth conditions, likely because transient stalling events of ribosomes occur naturally during unperturbed translation.

**Figure 4.**
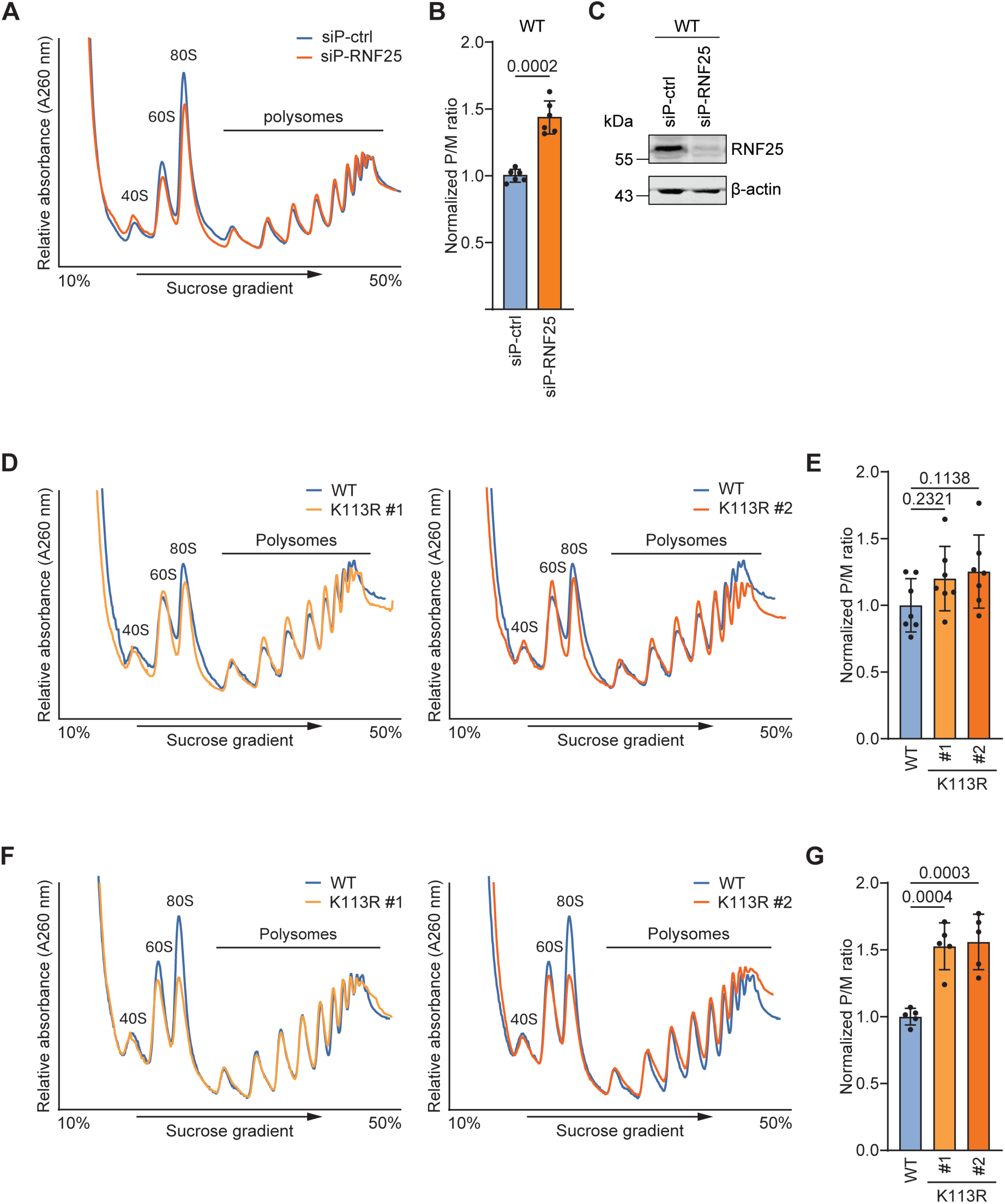
Ubiquitination of RPS27A/eS31 is needed to support translation. **(A)** Analysis of polysome profiles upon RNF25 knockdown. RNF25 was depleted from HeLa cells by treatment with siPOOLs for 72 h. Cell extracts were separated by sedimentation in 10%-50% sucrose density gradients and profiles were visualized at 260 nm. **(B)** Quantification of polysome/monosome (P/M) area ratios in profiles from (A), normalized to control profiles (N = 4, mean ± SD, unpaired two-tailed Student’s t-test, p-value = 0.0002). **(C)** Validation of knockdown experiments from (A). The efficiency of downregulation was analysed by immunoblotting of cell extracts with the indicated antibodies. **(D)** Analysis of polysome profiles upon inactivation of the RNF25 pathway. Cell extracts from parental HeLa WT cells or gene-edited *RPS27A* K113R cells were separated by sedimentation in 10%-50% sucrose density gradients and profiles were visualized at 260 nm. **(E)** Quantification of polysome/monosome (P/M) area ratios in profiles from (D), normalized to WT profiles (N = 7, mean ± SD, one-way ANOVA and post hoc Dunnett’s test, p-values are indicated). **(F)** Analysis of polysome profiles of HeLa WT and RPS27A K113R cells upon ISRIB treatment. Parental HeLa WT cells or gene-edited *RPS27A* K113R cells were treated with 1.1 µM ISRIB for 6 h. Cell extracts were separated by sedimentation in 10%-50% sucrose density gradients and profiles were visualized at 260 nm. **(G)** Quantification of polysome/monosome (P/M) area ratios in profiles from (F), normalized to WT profiles (N = 5, mean ± SD, one-way ANOVA and post hoc Dunnett’s test, p-values are indicated).

To examine the functional importance of the RNF25-mediated ubiquitination of RPS27A/eS31 in this regard, we utilized genome-edited HeLa cells harbouring a K113R substitution in RPS27A/eS31, which prevents its RNF25-dependent ubiquitination (Fig S3A, S3B). These cells are therefore deficient in the RNF25 pathway, evident by the lack of eRF1 degradation upon treatment of cells with SRI-41315 (Fig S3B). Curiously though, there was no significant difference in the P/M ratio in lysates of K113R and WT cells growing under optimal conditions, although the P/M ratio was slightly elevated (Fig 4D, 4E). We reasoned that the difference to the RNF25 knockdown experiments might be explained by the possibility that upon long-term inactivation of the RNF25 pathway, as in the K113R mutant cells, other pathways take over to resolve stalled ribosomes. Intriguingly, when cultivating K113R cells in the presence of ISRIB, which activates translation by reducing the inhibitory effect of phospho-eIF2α and thereby antagonising the ISR, we could see a substantial increase in the P/M ratio compared to WT cells (Fig 4F, 4G). These data further support the notion that activation of the RNF25 pathway is needed not only to deal with collisions caused by external stimuli, but also to resolve transient ribosome collisions occurring naturally under optimal growth conditions.

### Level of ubiquitinated RPS27A/eS31 increases in the absence of GCN2 and DRG2/RWDD1

Our data thus far indicate that RNF25-dependent ubiquitination of RPS27A/eS31 is dependent on the presence of GCN1 and is induced upon amino acid starvation (Fig 1E-G and 3B-F). GCN1 is not only involved in the RNF25 pathway, but it is also a part of the ISR, which can be induced by multiple triggers including amino acid starvation. In the ISR pathway, GCN1 recognizes collided ribosomes and recruits GCN2, which is subsequently activated and autophosphorylated (Darnell *et al*., 2018; Hinnebusch, 1994, 2005; Marton *et al*., 1993; Masson, 2019; Zhou *et al*., 2025). Thus, both GCN2 and RNF25 rely on the recognition of collided ribosomes by GCN1 for their activation. There are at least two possible scenarios to reconcile the functional relationship between these factors: either GCN2 and RNF25 belong to the same pathway, or they are parts of two independent parallel pathways, both depending on GCN1.

To examine whether RNF25 is involved in the activation of the ISR, we compared levels of phospho-GCN2 in WT and K113R cells upon amino acid starvation (Fig S4A, S4B). Immunoblot analysis revealed comparable levels of phospho-GCN2 between the different cell lines, indicating that ubiquitination of RPS27A/eS31 by RNF25 is not required for the activation of GCN2 as part of the ISR and that the two pathways diverge downstream of GCN1.

To test the idea that the ISR and RNF25 pathways co-exist as two parallel pathways, we generated *USP16/GCN2* DKO cell lines from parental HeLa *USP16* KO cells (Fig S4C, S4D). Interestingly, the levels of ubiquitinated RPS27A/eS31 increased in the absence of GCN2 already in non-starved conditions (Fig 5A, 5B), and there was no further increase in the ubiquitination levels upon amino acid starvation. This indicates that in the absence of GCN2, the RNF25 pathway is already fully activated by naturally occurring translational stalling events and supports the hypothesis that RNF25 and GCN2 represent two independent pathways downstream of GCN1, guarding cells against ribosome collisions.

**Figure 5.**
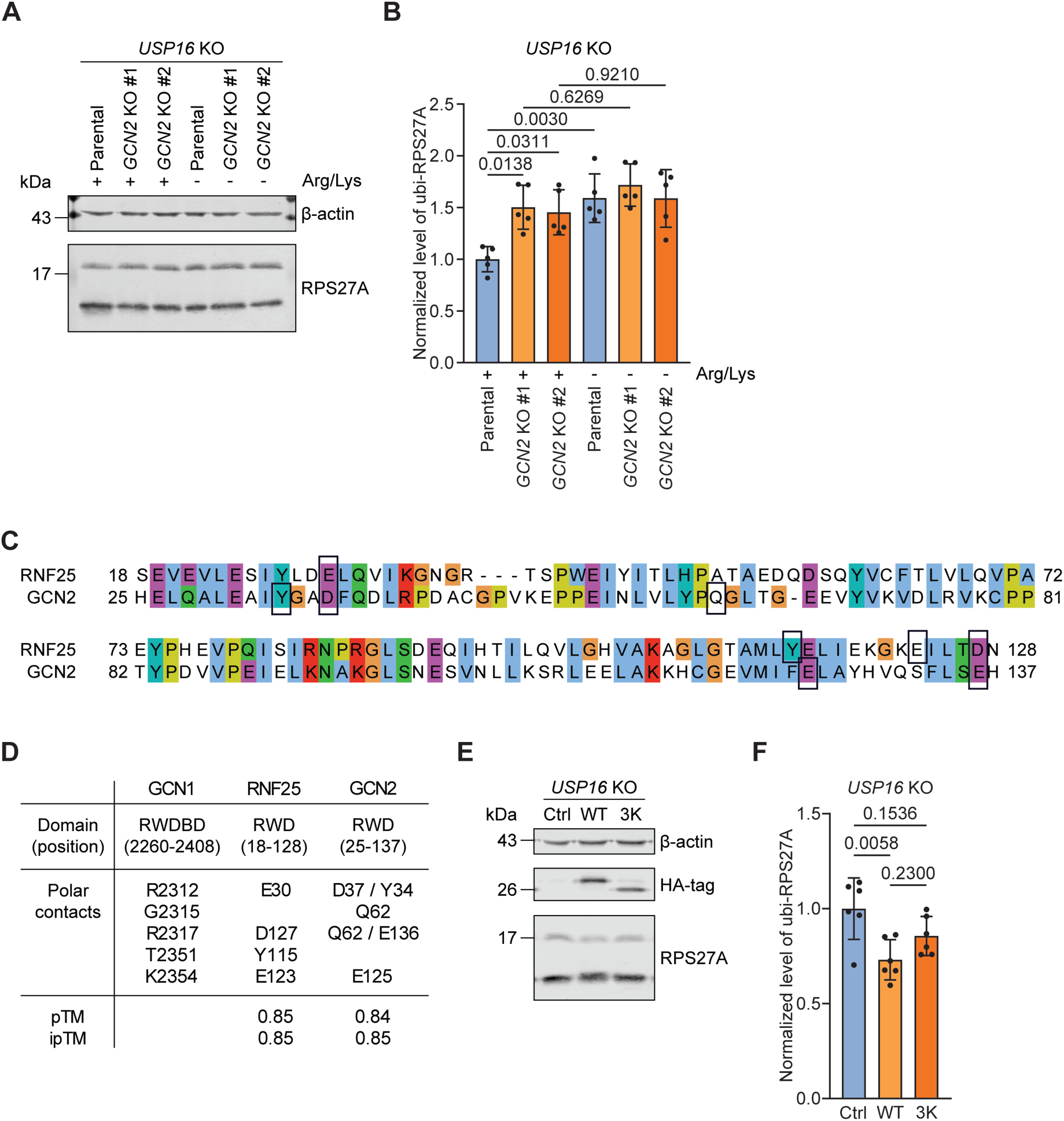
Level of ubiquitinated RPS27A/eS31 increases in the absence of GCN2. **(A)** Analysis of RPS27A/eS31 ubiquitination in HeLa *USP16* KO cells upon deletion of *GCN2*. The *GCN2* gene was deleted from *USP16* KO cells by CRISPR/Cas9. The parental *USP16* KO cells and two clones of *USP16*/GCN2 double knockout (DKO) cells were cultivated for 6 h in the presence (+) or absence (-) of arginine and lysine. Cell extracts were analysed by immunoblotting with the indicated antibodies. **(B)** Quantification of levels of ubiquitinated RPS27A/eS31 in cell lysates from (A), expressed as the ratio between ubi-RPS27A/eS31 and total RPS27A/eS31 (ubi-RPS27A/eS31 + unmodified RPS27A/eS31) and normalized to samples from non-starved parental *USP16* KO cells (N = 5, mean ± SD, one-way ANOVA and post hoc Tukey’s test, p-values are indicated). **(C)** Sequence alignment of RWD domains of RNF25 (aa residues 18-128) and GCN2 (aa residues 25-137). Sequences were visualised in Jalview and aligned with Clustal W. Residues predicted by AlphaFold 3 web server to interact with RWDBD domain of GCN1 are indicated with rectangles. Residues are colored according to the Clustal X color scheme: blue – hydrophobic, red – positive charge, magenta – negative charge, green – polar, orange – glycine, yellow – proline, cyan – aromatic. **(D)** Prediction of interaction between RWDBD domain of GCN1 and RWD domains of RNF25 and GCN1 using AlphaFold 3. Protein domains as well as positions of respective residues used for the modelling are indicated. Respective pTM and ipTM scores show high-quality predictions (> 0.8) for both the overall protein structure (pTM) and the protein-protein complex (ipTM). Indicated polar contacts were analysed with PyMol. **(E)** Analysis of RPS27A/eS31 ubiquitination in HeLa *USP16* KO cells upon transient overexpression of the indicated GCN2 fragments containing an N-terminally StStHA-tagged RWD domain (aa 1-145). The effect of WT and 3K mutant (D37K, E125K and E136K substitutions) RWD domains was compared to control conditions (transfection of empty vector). Cell extracts were analysed by immunoblotting with the indicated antibodies. **(F)** Quantification of levels of ubiquitinated RPS27A/eS31 in cell lysates from (E), expressed as the ratio between ubi-RPS27A/eS31 and total RPS27A/eS31 (ubi-RPS27A/eS31 + unmodified RPS27A/eS31) and normalized to control samples (N = 6, mean ± SD, one-way ANOVA and post hoc Tukey’s test, p-values are indicated).

Both RNF25 and GCN2 possess RWD domains that can bind the RWDBD domain of GCN1, raising the possibility that RNF25 and GCN2 compete for GCN1 binding via their RWD domains. Comparative analysis of the respective RWD domains with ClustalW as well as AlphaFold prediction of interactions between these domains and the RWDBD domain of GCN1 suggest that binding occurs using a similar interaction interface (Fig 5C, 5D). In both cases, polar interactions involved GCN1 residues R2312, R2317 and K2354, with R2312 forming contacts with the positionally conserved residues E30 of RNF25 or D37 of GCN2 (Fig 5C, 5D). This model agrees with previous observations describing that a R2312A substitution in GCN1 prevents binding and activation of GCN2 (Sattlegger & Hinnebusch, 2000; Zhou *et al*., 2025). To test the idea that the RWD domains of RNF25 and GCN2 compete for GCN1 binding, we overexpressed the wild-type RWD domain of GCN2 in HeLa *USP16* KO cells, as well as a mutant version harbouring D37K, E125K and E136K substitutions (3K mutant), which potentially has a hampered interaction with GCN1. The levels of ubiquitinated RPS27A/eS31 decreased about 1.4-fold upon expression of wild-type RWD domain of GCN2 (Fig 5E, 5F), supporting the idea that this domain competes with RNF25 for GCN1 binding. Cells expressing the 3K mutant had intermediate levels of ubiquitinated RPS27A/eS31, indicating that the interface mutations weaken GCN1 interaction. Altogether, these data support the idea that the RWD domains of RNF25 and GCN2 compete for GCN1 binding.

It is important to note that two further proteins, RWDD1 and Impact, contain RWD domains capable of GCN1 binding (Ishikawa *et al*, 2013; Pereira *et al*, 2005; Pochopien *et al*., 2021; Waller *et al*, 2012; Wout *et al*, 2009). Yeast homologs of RWDD1 and its cofactor DRG2, with the respective names Gir2 and Rbg2, have been previously identified in a cryo-EM model of collided ribosomes, where both proteins reside in the A-site factor binding region of the leading ribosome, and Gir2 mediates an interaction between the Rbg2/Gir2 complex and Gcn1 (Pochopien *et al*., 2021). To test whether the human DRG2/RWDD1 complex represents another pathway competing with RNF25, we generated *USP16/DRG2* DKO cells using parental HeLa *USP16* KO (Fig S5A). As expected (Ishikawa *et al*, 2009), these cells exhibited decreased levels of RWDD1 upon depletion of its cofactor DRG2, reflecting the degradation of excess RWDD1 protein when formation of the DRG2/RWDD1 complex is prevented (Fig S5B, S5C). In turn, the levels of ubiquitinated RPS27A/eS31 increased in the absence of DRG2 (Fig S5B, S5D), which might be caused by reduced competition between RWDD1 and RNF25, and a consequent inability to resolve ribosome stalling through the DRG2 pathway. Altogether, these results suggest that GCN1-dependent quality control of protein synthesis has evolved as a multifaceted process involving several parallel downstream pathways.

## DISCUSSION

Recent studies have identified a novel translation-dependent quality control pathway involving the E3 ligase RNF25, which ubiquitinates RPS27A/eS31 (Gurzeler *et al*., 2023; Oltion *et al*., 2023). Here, we used ubiquitination levels of RPS27A/eS31 as an indicator of RNF25 activation to systematically investigate the triggers and mechanism of this pathway.

Our data indicate that ubiquitination of RPS27A/eS31 is induced by a variety of conditions that cause ribosome stalling. First, we established that besides ternatin, the presence of other translation inhibitors, such as emetine, didemnin B, anisomycin or MMS, leads to the activation of RNF25. Interestingly, these drugs have different modes of action, not always leading to trapped translation factors in the ribosomal A-site. Didemnin B acts similarly to ternatin by locking eEF1A in the A-site of the ribosome (Juette *et al*., 2022). Emetine binds to the E-site of the 40S subunit and prevents translocation, leaving eEF2 bound to the ribosomal A-site (Wong *et al*., 2014; Zhou *et al*., 2025). Anisomycin binds at the peptidyl transferase centre of the 60S subunit and inhibits peptide bond formation. In the presence of anisomycin, incoming aminoacyl-tRNA can still undergo the codon-anticodon pairing step, but the correct placement of the aminoacyl-3’-CAA end of the tRNA is abolished. This stalls ribosomes in the pre-peptide bond formation conformation with a partially accommodated tRNA (Garreau de Loubresse *et al*., 2014; Wu *et al*, 2019a). MMS is a nucleic acid damaging reagent that induces RNA methylation. Presence of bulky methyl groups on mRNAs was suggested to prevent correct aminoacyl-tRNA accommodation, leaving ribosomes stalled with an empty A-site (Stoneley *et al*., 2022; Thomas *et al*., 2020; Yan & Zaher, 2021). Importantly, we could also identify amino acid starvation as a physiological trigger that activates the RNF25 pathway, leading to the ubiquitination of RPS27A/eS31. Under starvation conditions, ribosome stalling is caused by the lack of aminoacyl-tRNA. However, uncharged tRNA might still bind to the A-site of the ribosome and base-pair with cognate codons with the help of the GCN2 kinase, which contributes to the activation of GCN2 on the ribosome (Zhou *et al*., 2025). Thus, our analyses demonstrate that a variety of triggers leading to diverse ribosomal states can activate RNF25, suggesting that activation of this pathway does not *per se* require the presence of trapped protein factors in the ribosomal A-site but is a more general response to ribosome collisions.

Ribosomal collisions are recognized by several sensors, namely EDF1, ZNF598, GCN1 and ZAKα. We have previously excluded a role for ZNF598 in RPS27A/eS31 ubiquitination (Montellese *et al*., 2020). Here, we investigated whether ubiquitination of RPS27A/eS31 depends on the presence of EDF1, GCN1 or ZAKα. Two of these sensors – EDF1 and GCN1 – are involved in the activation of the ISR pathway (Kim *et al*., 2024; Marton *et al*., 1993; Pochopien *et al*., 2021), while ZAKα is involved in the RSR pathway (Huso *et al*., 2026; Sinha *et al*., 2024; Vind *et al*., 2020). Our analysis demonstrated that ubiquitination of RPS27A/eS31 does not depend on the presence of EDF1 or ZAKα, but on GCN1, in agreement with earlier observations (Gurzeler *et al*., 2023; Oltion *et al*., 2023). A previous structural study of collided disomes in yeast has identified the C-terminal part of Gcn1 (with its RWD domain) in vicinity of yeast Rps31/eS31 (Pochopien *et al*., 2021), suggesting that also on mammalian disomes, GCN1 would be ideally positioned to guide RNF25 to RPS27A/eS31 for its ubiquitination.

Intriguingly, our analysis showed that the RNF25 pathway is needed to support translation already under optimal growth conditions, as inactivation of this pathway leads to an increased polysome/monosome ratio, hinting at the general presence of low levels of unresolved ribosome collisions. This indicates that the RNF25 pathway is active to resolve collisions caused not only by translational stress, but also transient events naturally occurring during active translation. Interestingly, a recent study using *in situ* cryo-electron tomography indeed detected GCN1 on 2% of collided ribosomes in untreated cells and at 6% of disomes upon anisomycin treatment, whereas it was excluded from ribosomes in compact and persistent collisions (Fedry *et al*, 2024). It has also been reported that transient ribosome collisions caused by nonoptimal codons or in 3’ UTR regions are recognized by GCN1 (Muller *et al*, 2023). Moreover, we previously showed that G418, a drug that induces stop codon readthrough, induces ubiquitination of RPS27A/eS31 (Montellese *et al*., 2020), suggesting that the RNF25 pathway is involved in the resolution of collisions in the 3’UTR of mRNAs. Since activity of the RNF25 pathway depends on the presence of GCN1, it is possible that GCN1 and RNF25 act in tandem to recognize “loose” collisions produced by transient stalling events.

GCN1 is an integral part of the ISR pathway, where it participates in the activation of GCN2 on ribosomes upon amino acid starvation. Active GCN2 inhibits translation initiation by phosphorylating the α subunit of eIF2. Surprisingly, we found that the level of ubiquitinated RPS27A/eS31 increases in the absence of GCN2 already in non-starved conditions, hinting at a competition between GCN2 and RNF25. Both proteins possess structurally similar RWD domains that can bind the RWDBD domain of GCN1, suggestive of a mechanism where competition of RNF25 and GCN2 is at least partially mediated by these domains. In support of this idea, we observed that overexpression of the RWD domain of GCN2 leads to a decreased level of ubiquitinated RPS27A/eS31. While this manuscript was in preparation, two other studies described competitive relations between RNF25 and GCN2, where RNF25 protects cells from the GCN2-dependent activation of ISR upon RNA damage (Seidel *et al*, 2026; Zhao *et al*, 2026). In turn, our data support a broader model according to which such competition occurs not only in response to RNA damage (MMS treatment, Fig 1B-D), but also during amino acid starvation as well as upon transient stalling in non-starved conditions (Fig 5A, 5B).

Further extending the complexity of the regulatory network, our data indicates that the DRG2/RWDD1 complex represents another GCN1-dependent pathway that competes with RNF25. To date, little is known about how the DRG2/RWDD1 pathway intersects with other RQC factors. A recent cryo-EM structure of yeast GCN1-bound disomes has visualised the yeast Rbg2/Gir2 complex (homologs of DRG2/RWDD1) in the A-site factor binding region of the ribosome, where Rbg2 directly interacts with the A-site tRNA and Gir2 mediates interaction of the complex with the RWDBD domain of Gcn1 (Pochopien *et al*., 2021). Based on this structure it has been suggested that Rbg2 stabilizes the accommodated A-site tRNA and promotes efficient translation upon ribosome pausing (Pochopien *et al*., 2021). Moreover, binding of the Rbg2/Gir2 complex has been shown to inhibit ISR activation by preventing the recruitment of Gcn2 (Ishikawa *et al*., 2013). Therefore, it is possible that in human cells the RNF25 and DRG2/RWDD1 pathways have overlapping functions. RNF25 may act as a general sensor of stalled ribosomes, recognizing ribosomes with empty, t-RNA and/or factor occupied A-sites. DRG2/RWDD1, in contrast, seems to specifically recognises stalled ribosomes that lack a protein factor but have tRNA bound in the A-site. However, more work must be done to unravel the relationship between these two pathways.

Collectively, our work revealed that (i) the RNF25 pathway can be activated by various triggers that affect A-site occupancy in distinct ways, (ii) loss of RNF25 increases the polysome/monosome ratio in untreated cells, suggesting a role in clearance of collisions that occur naturally during translation, (iii) GCN1 serves as a central factor of a regulatory network at which GCN2 and DRG2/RWDD1 compete with the RNF25 pathway. Based on these observations, we propose a model in which the RNF25 pathway is activated as a general response to a variety of ribosome collisions (low doses of translation inhibitors, RNA damage, amino acid starvation, transient collision events) (Fig 6). For the activation of this pathway, such ribosome collisions are first recognized by GCN1, which subsequently recruits RNF25 via its RWD domain. As a result, RNF25 ubiquitinates RPS27A/eS31, which in turn induces the resolution of stalled ribosomes. Initially, RNF25, which is likely only present in very low amounts (23’300 molecules per HeLa cell (Itzhak *et al*, 2016)), is recruited to resolve such transient stalling events. As a parallel pathway, DRG2/RWDD1 recognizes specific stalling events where ribosomes slow down and have a vacant A-site factor binding region. Once collisions increase above a certain threshold or persist, GCN2 is activated, likely with overall increasing amount of GCN1 on collided ribosomes, leading to the activation of the IRS pathway and inhibition of translation.

**Figure 6.**
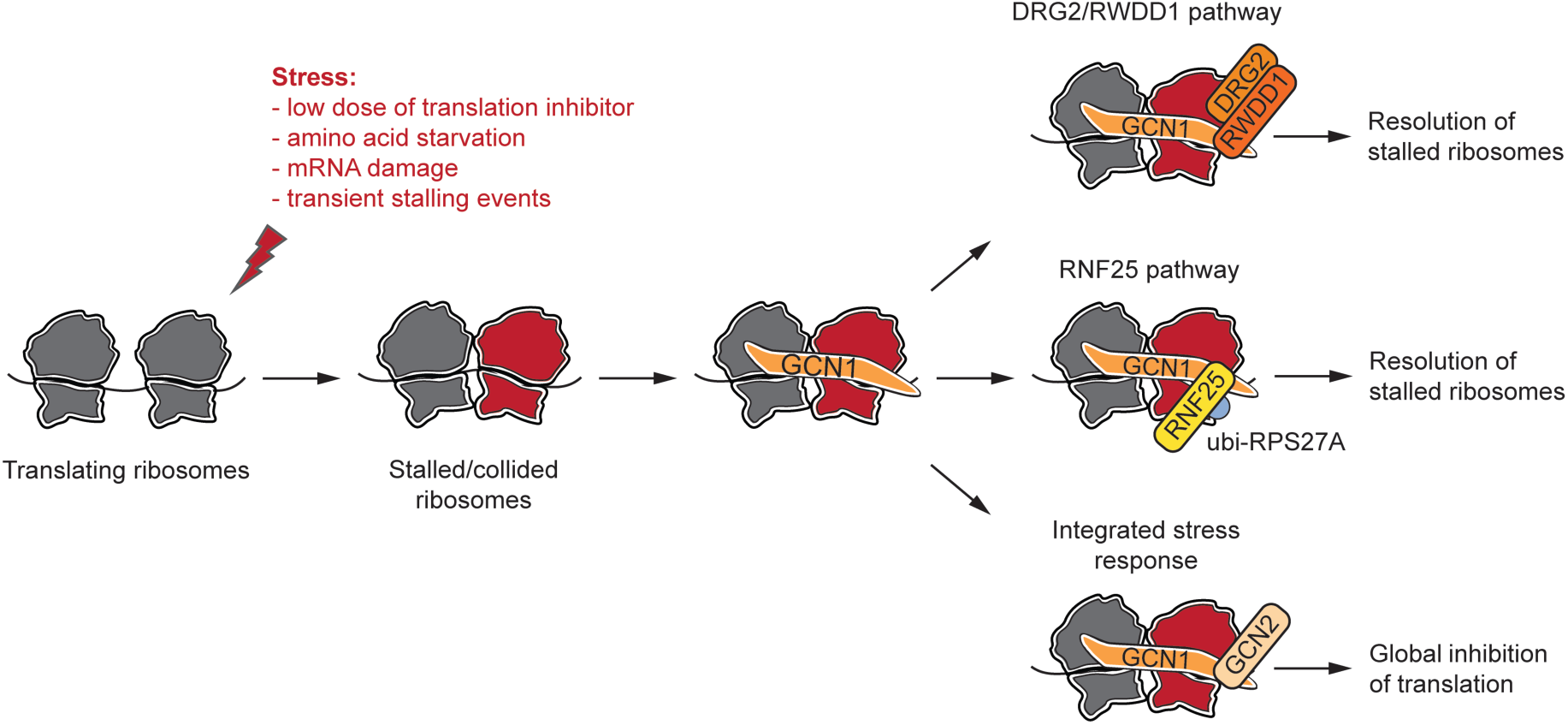
Model for the activation of the RNF25 pathway. Upon conditions that induce ribosome stalling (such as low doses of translation inhibitors, mRNA damage or amino acid starvation) as well as upon natural transient stalling events, collided ribosomes are recognized by GCN1. Following GCN1 binding, RNF25 is recruited onto ribosomes through the interaction between its RWD domain and the RWDBD domain of GCN1, resulting in the ubiquitination of RPS27A/eS31 and subsequent resolution of stalled ribosomes. Alternatively, the DRG2/RWDD1 complex can also interact with GCN1 and engage in resolution of stalled ribosomes that have no protein factor in the A-site factor binding region, but a tRNA bound in the A-site. With an increased number of collisions or more persistent collisions, GCN1 recruits GCN2 through interaction with its RWD domain, which induces activation of the ISR pathway and inhibition of translation.

## MATERIALS AND METHODS

### Cell lines, culturing conditions, and treatments

HeLa cells were cultured in DMEM supplemented with 10% FCS and 100 µg/ml penicillin/streptomycin at 37°C and 5% CO_2_. HeLa K WT cells and HeLa K *USP16* KO were described previously (Montellese *et al*., 2020). Where indicated, cells were treated with following compounds: anisomycin (Sigma, A9789), emetine (Sigma, E2375), didemnin B (Sigma, SML4022), ternatin (Santa Cruz, sc-391653), methyl methanesulfonate (Sigma, 129925), SRI-41315 (Sigma, SML3312), ISRIB (Sigma, SML0843).

In amino acid starvation experiments, cells were grown in DMEM for SILAC (Thermo Scientific, 88364) supplemented with 10% dialyzed FCS (Gibco, A3382001), 100 µg/ml penicillin/streptomycin, 84 µg/ml arginine and 146 µg/ml lysine. To induce amino acid starvation, arginine and/or lysine were omitted from the media where indicated.

### RNA interference

HeLa cells growing on either 6-well or 10 cm plates were transfected with 5 nM siPOOLs (siTOOLs, (Hannus *et al*, 2014)) in Opti-MEM (Gibco, 31985047) using INTERFERin (Polyplus, 101000028). Medium was exchanged on the next day.

### Molecular cloning and transfection of RWD domain-encoding constructs

DNA fragments encoding for the RWD domain of human GCN2 (aa residues 1-147, WT and 3K mutant) were ordered from TWIST Bioscience and inserted into pcDNA/FRT/TO vector (Invitrogen) with N-terminal StStHA-tag using KpnI and XhoI cloning sites.

HeLa K *USP16* KO cells growing on 6-well plates were transfected with the RWD domain encoding vectors using the jetPrime reagent (Sartorius, 101000046). Media was exchanged 6 h post-transfection, and cells were harvested 48 h post-transfection.

### Generation of knockout cell lines

Double knockout (DKO) HeLa cell lines (*USP16/RNF25* DKO, *USP16/GCN1* DKO, *USP16/GCN2* DKO and *USP16/DRG2* DKO) were generated using a CRISPR-Cas9 genome editing system. The following gRNA sequences were used in this study: RNF25 exon 3: 5’ AAAGTGATGTAGATCTCCCA 3’ (Oltion *et al*., 2023) GCN1 exon 14: 5’ AAACCTCCACATCTGCGGTG 3’ (Doench *et al*, 2016) GCN2 exon 3: 5’ ACTGGCCAAGAAACACTGTG 3’ (Doench *et al*., 2016) DRG2 exon 1: 5’ GCTCGGACACAGAAGAACAA 3’ (Doench *et al*., 2016) gRNA sequences were inserted into the BsaI site of the pC2P vector containing hCas9 and a puromycin resistance cassette (Welte *et al*, 2019). HeLa K *USP16* KO cells growing on 6-well plates were transfected with the resulting vector using the jetPrime reagent (Sartorius, 101000046). The next day, cells were re-seeded onto 10 cm plates. After 24 h, and selection was started with 1 µg/ml of puromycin for 2 days. After selection, individual clones were expanded, and the knockouts were verified by Sanger sequencing and immunoblotting. Sequencing results were decomposed either manually or using TIDE web service (Brinkman *et al*, 2014).

### Prime editing

HeLa K cells harbouring the base substitutions at the endogenous RPS27A/eS31 loci for expression of the K113R mutant were generated using prime editing (Doman *et al*, 2022). Specifically, the sequence 5’-GAATGGC**AAA**ATT-3’ located at position 55235436-55235448 on the positive strand of chromosome 2 was substituted to a sequence 5’-GAA<u>CGGC**CG**</u>**A**ATT-3’. This resulted in the substitution of AAA®CGA (Lys®Arg, bold) and introduction of a silent EagI digestion site (underlined). epegRNA targeting RPS27A/eS31 was designed using PrimeDesign tool (Hsu *et al*, 2021), and consisted of the following parts:

Spacer: 5’ CAGAAGGGCACTCTCGACGA 3’

Scaffold: 5’ CTAGAAATAGCAAGTTAAAATAAGGCTAGTCCGTTATCAACTTGAAAAAGTGGCACCGAGTCG 3’

Extension (41 nt RTT and 12 nt PBS): 5’ GCTTTTCAGGTGGATGAGAACGGCCGAATTAGTCGGCTTCGTCGAGAGTGCCC 3’

Spacer, scaffold and extension of epegRNA were assembled using Golden Gate cloning and inserted into the BsaI site of the pU6-tevopreq1-GG-acceptor vector, which was a gift from David Liu (Addgene plasmid # 174038) (Nelson *et al*, 2022). PEmax prime editor was expressed from the pCMV-PEmax vector, encoding for SpCas9 and MMLV-RT (a gift from David Liu, Addgene plasmid # 174820) (Chen *et al*, 2021). pIRESpuro encoding for a puromycin resistance cassette was used for selection of cells with puromycin.

HeLa K WT cells growing on 6-well plates were transfected with pIRESpuro, PEmax and epegRNA vectors at a ratio 0.1 ug : 1 ug : 1.2 ug using jetPrime. Next day, cells were re-seeded onto 10 cm plates. After 24 h, cells were selected with 1 µg/ml puromycin for 2 days. After selection, individual clones were picked and expanded. To verify the edits, genomic DNA was extracted with phenol/chloroform/IAA (Applichem, A0889.0500) and analyzed by digestion of PCR products with EagI and Sanger sequencing.

### Immunoblot analysis and antibodies

Cells growing on 6-well plates were washed once with ice-cold 1x PBS and collected by scraping in 120 µl of lysis buffer (50 mM HEPES-KOH pH 7.5, 150 mM NaCl, 1% NP-40, 5 mM N-ethylmaleimide, 1 ug/ml pepstatin, 10 ug/ml leupeptin, 10 ug/ml aprotinin). In case of analysis of phosphorylated proteins, lysis buffer was additionally supplemented with 0.5 mM NaF and 0.1 mM NaVO_4_. Cells were incubated on ice for 5 min with occasional vortexing, and the resulting lysates were cleared by centrifugation (16,000 x g, 10 min, 4°C). Protein concentration was measured with the Bradford method using the Bio-Rad Protein Assay Dye Reagent Concentrate (Bio-Rad, 5000006), and equal amounts of protein were supplemented with SDS loading buffer. Samples were denatured for 5 min at 95°C and resolved on Tris-glycine SDS-polyacrylamide gels. Proteins were transferred to nitrocellulose membranes by semi-dry blotting. Membranes were blocked in 4% milk in PBST and incubated overnight at 4°C with primary antibodies diluted in 4% milk/PBST. Next, membranes were washed 3x 10 min in PBST and incubated with secondary antibodies diluted in 4% milk/PBST at room temperature for 1 h. Membranes were washed for 3x 10 min in PBST, and signals were detected using an Odyssey (LI-COR) imaging system. Signal intensities were measured with Fiji (Schindelin *et al*, 2012). Statistical analysis was done in GraphPad Prism 10.

The antibody against RPS27A/eS31 was described previously (Montellese *et al*., 2020). Other primary antibodies that were used in this study are: anti-β-actin (1:2000, Santa Cruz, sc-47778), anti-eRF1 (1:1000, Santa Cruz, sc-365686), anti-RNF25 (1:500, Santa Cruz, sc-398749), anti-GCN1 (1:1000, Bethyl, A301-843A), anti-GCN2 (1:1000, Abcam, ab134053), anti-phospho-GCN2 (Thr899) (1:500, Cell Signaling, 94668), anti-EDF1 (1:1000, Abcam, ab174651), anti-ZAKα (1:1000, Bethyl, A301-993A), anti-HA (1:1000, Enzo, ENZ-ABS120-0200). Antibodies against human DRG2 and RWDD1 were raised in rabbits against purified recombinant His_6_-tagged proteins expressed in *E. coli*. Secondary antibodies used in this study were: goat anti-mouse IgG (H+L) Alexa Fluor^TM^ 680 (1:10000, Invitrogen, A21058), goat anti-rabbit IgG (H+L) Alexa Fluor^TM^ Plus 800 (1:10000, Invitrogen, A32735).

### Sucrose density gradients

HeLa cells were grown on 10 cm plates and treated with 100 µg/ml cycloheximide (Sigma, C7698) for 10 min prior to harvesting. Cells were washed once with ice-cold 1x PBS containing 100 µg/ml cycloheximide and collected by scraping in 400 µl of lysis buffer (50 mM HEPES-KOH pH 7.5, 100 mM KAc, 10 mM MgAc_2_, 0.5 % NP-40, 1 mM DTT, 100 µg/ml cycloheximide, 1 µg/ml pepstatin, 10 µg/ml leupeptin, 10 µg/ml aprotinin, 40 U/ml RiboLock RNase inhibitor (Thermo Fisher, EO0381). Collected cells were incubated on ice for 5 min and the resulting lysates were cleared by centrifugation (16,000 x g, 10 min, 4°C). The A260 of lysates was measured with NanoDrop. Equal amounts of A260 units were loaded onto 10%-50% sucrose gradients produced in gradient buffer (50 mM HEPES-KOH pH 7.5, 100 mM KAc, 10 mM MgAc_2_, 1 mM DTT, 100 µg/ml cycloheximide). Lysates were centrifuged in a SW41 rotor for 2 h at 39,000 rpm, and the A260 profiles of the gradients were monitored using a BioComp fractionator at the A260 during gradient fractionation. Areas under the monosome and polysome peaks were measured with Fiji (Schindelin *et al*., 2012), and statistical analysis was done in GraphPad Prism 10.

### Visualization and modelling

Protein sequences were visualized in Jalview and aligned using the Clustal W (Procter *et al*, 2021; Waterhouse *et al*, 2009). Protein interactions were predicted using the AlphaFold 3 web server (Abramson *et al*, 2024).

## ACKNOWLEDGEMENTS

We thank Bianka Horváth for critical reading of the manuscript, members of the Kutay lab for helpful discussion, J. uit de Bos for advice on prime editing, I. Lazarevic and B. Widmann for generation of anti-DRG2 and anti-RWDD1 antibodies, and C. Montellese for initial work on RPS27A gene-editing. We are grateful to our colleagues C.M.T. Spahn and F. Wiechert (Universitätsmedizin Berlin) as well as O. Mühlemann and W. Teodorowicz (University of Bern) for helpful discussions and collaboration on related aspects.

## FUNDING

This work was supported by the Swiss National Science Foundation (SNSF) in the framework of the NCCR “RNA and disease” (51NF40-205601) and by the SNSF Advanced Grant TMAG-3_209245 to U.K..

## DATA AVAILABILITY

Our study includes no data deposited in public repositories.

## DISCLOSURE AND COMPETING INTEREST STATEMENT

The authors declare no conflict of interest.

## Notes

### Competing Interest Statement

The authors have declared no competing interest.

