## Supplemental Material for "RNF25 is activated as a response to amino acid starvation-induced ribosome collisions in competition with GCN2"

### SUPPLEMENTAL MATERIALS

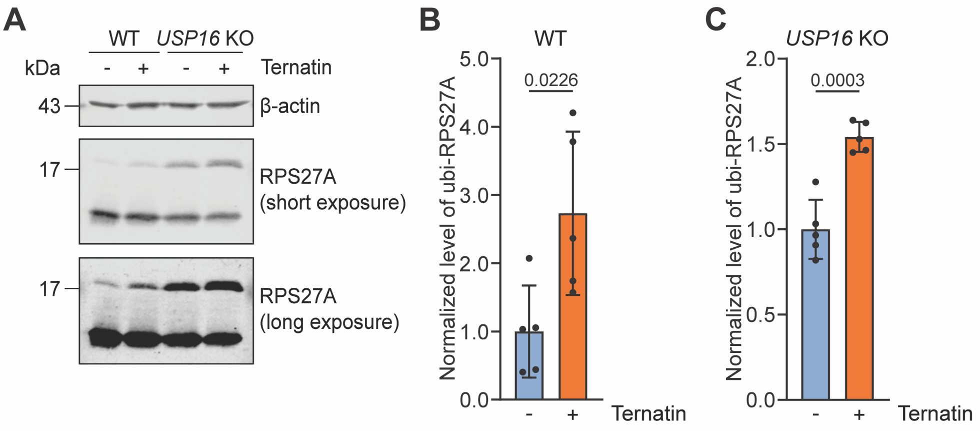


**Supplemental Figure S1. Ubiquitination of RPS27A/eS31 is induced by ternatin treatment in both WT and *USP16* KO cells.**

**(A**) The eEF1A inhibitor ternatin induces RPS27A/eS31 ubiquitination. HeLa WT and *USP16* KO cells were cultivated for 4 h in the absence or presence of 50 nM ternatin. Cell extracts were analysed by immunoblotting with the indicated antibodies.

**(B, C)** Quantification of levels of ubiquitinated RPS27A/eS31 in HeLa WT (B) and *USP16* KO (C) lysates from (A), expressed as the ratio between ubi-RPS27A/eS31 and total RPS27A/eS31 (ubi-RPS27A/eS31 + unmodified RPS27A/eS31) and normalized to samples from untreated cells (N = 5, mean ± SD, unpaired two-tailed Student's t-test, p-values are indicated).

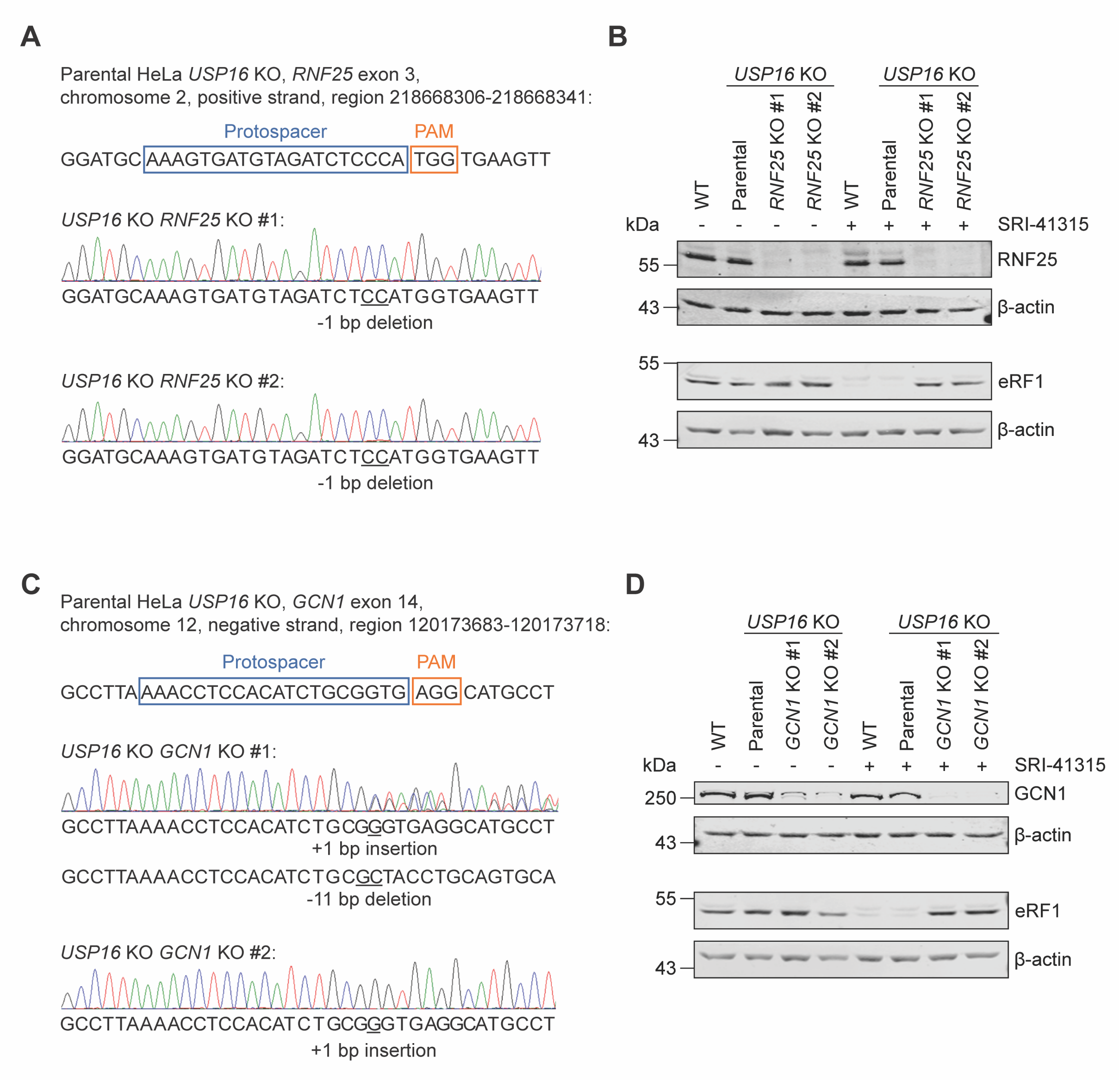


**Supplemental Figure S2. Validation of *USP16/RNF25* and *USP16/GCN1* DKO cell lines.**

**(A)** Validation of HeLa *USP16/RNF25* DKO cell lines, generated using the CRISPR/Cas9 system in parental *USP16* KO cells. At the top, the target region in the exon 3 of the *RNF25* gene in parental cells is represented. Positions of protospacer and PAM site are indicated. Below, Sanger sequencing chromatograms of the two generated clones are shown. Respective sequences after manual decomposition of chromatograms as well as introduced indel mutations are displayed.

**(B)** Immunoblot analysis of cell extracts from clones described in (A) using the indicated antibodies. To control for the expected inactivation of the eRF1-degradation pathway in the *USP16/RNF25* DKO clones, cells were cultivated in the absence (-) or presence (+) of 10 μM SRI-41315 for 20 h.

**(C)** Validation of HeLa *USP16/GCN1* DKO cell lines generated using the CRISPR/Cas9 system in parental *USP16* KO cells. At the top, the target region in the exon 14 (out of 54) of the *GCN1* gene in parental cells is represented. Positions of protospacer and PAM site are indicated. Below, Sanger sequencing chromatograms of the two generated clones are shown. Respective sequences after manual decomposition of chromatograms as well as introduced indel mutations are displayed.

**(D)** Immunoblot analysis of cell extracts from clones described in (C) using the indicated antibodies. *USP16/GCN1* DKO clones display low levels of residual full-length GCN1. To control for the expected inactivation of the eRF1-degradation pathway in DKO clones, cells were cultivated in the absence (-) or presence (+) of 10 µM of the eRF1 inhibitor SRI-41315 for 20 h.

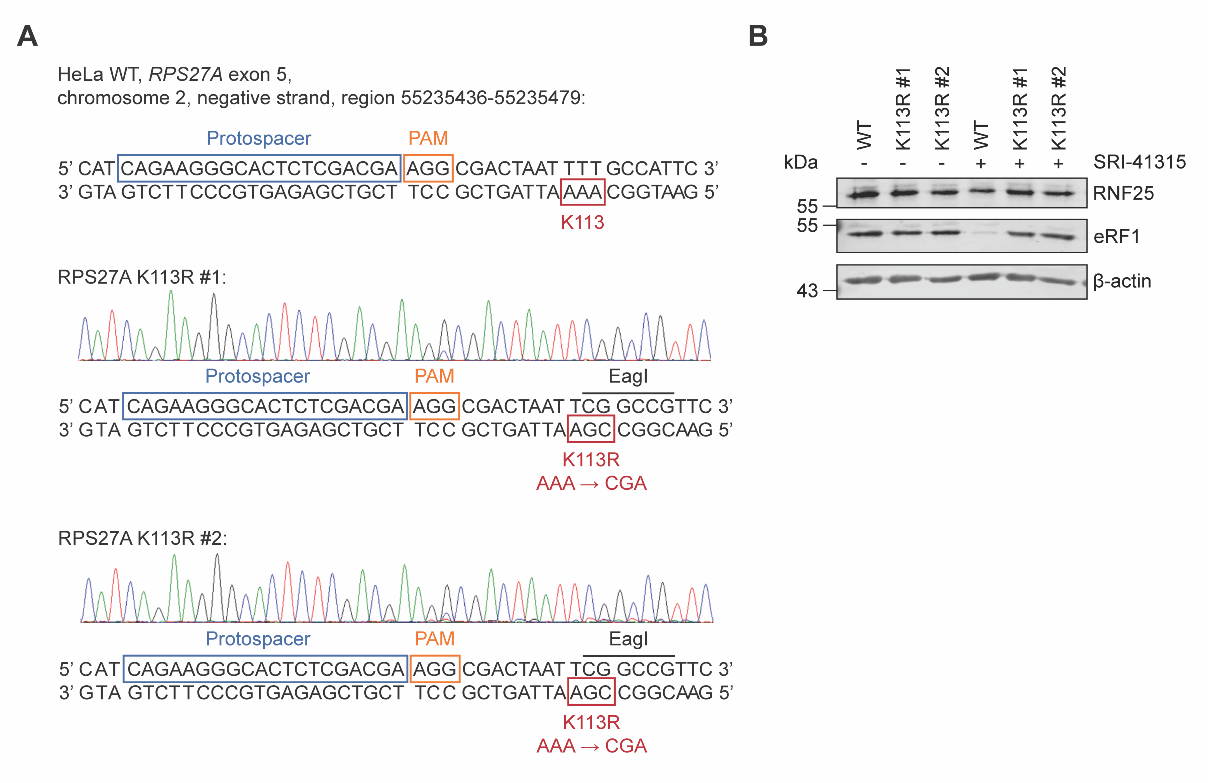


**Supplemental Figure S3. Validation of gene-edited *RPS27A* K113R cell lines.**

**(A)** Validation of gene-edited HeLa *RPS27A* K113R cell lines, generated using a prime-editing system in parental HeLa WT cells. At the top, the target region in exon 5 of the *RPS27A* gene in parental cells is represented. Positions of protospacer, PAM site and AAA codon encoding for K113 are indicated. Below, Sanger sequencing chromatograms of the two generated clones after prime editing. Respective sequences with K113R substitution (AAA → CGA) as well as the EagI site thereby introduced are displayed.

**(B)** Immunoblot analysis of cell extracts from clones described in (A). To control for the expected inactivation of the eRF1-degradation pathway in the generated K113R clones, cells were cultivated in the absence (-) or presence (+) of 10 µM SRI-41315 for 20 h.

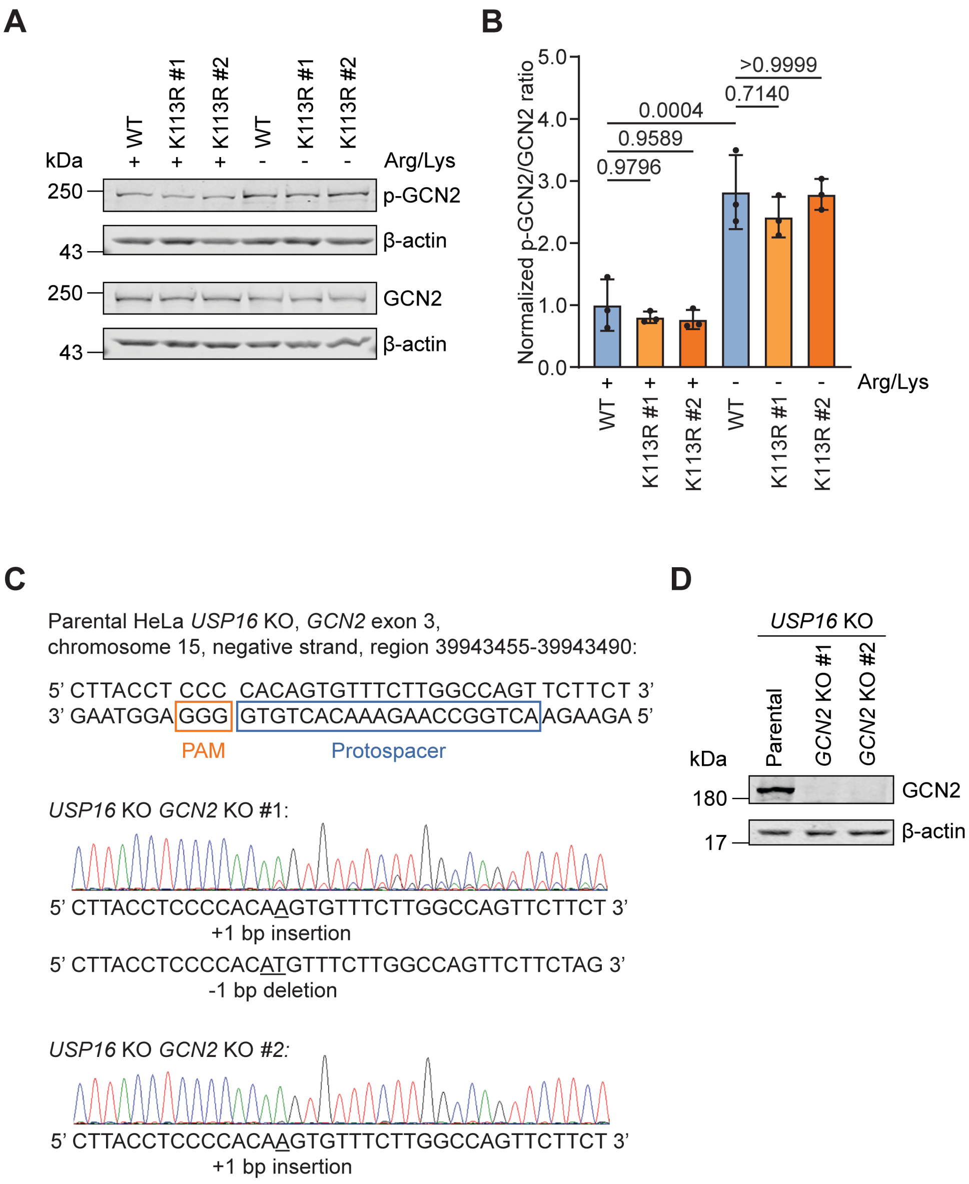


**Supplemental Figure S4. GCN2 activation upon amino acid starvation is independent of RNF25, and validation of *USP16/GCN2* double knockout cell lines.**

**(A)** Analysis of GCN2 phosphorylation upon inactivation of the RNF25 pathway. HeLa WT and K113R cells were cultivated for 6 h in the presence (+) or absence (-) of arginine and lysine. Cell extracts were analysed by immunoblotting with the indicated antibodies.

**(B)** Quantification of phosphorylated GCN2 in cell lysates from (A), expressed as the ratio between phosphorylated and unmodified GCN2, and normalized to samples from non-starved WT cells (N = 3, mean ± SD, one-way ANOVA and post hoc Tukey's test, p-values are indicated).

**(C**) Validation of HeLa *USP16/GCN2* DKO cell lines, generated using the CRISPR/Cas9 system in parental *USP16* KO cells. At the top, the target region in the exon 3 of the *GCN2* gene in parental cells is represented. Positions of protospacer and

PAM site are indicated. Below, Sanger sequencing chromatograms of the two generated clones are shown. Respective sequences after manual decomposition of chromatograms as well as introduced indel mutations are displayed.

**(D)** Immunoblot analysis of cell extracts from clones described in (C) using the indicated antibodies.

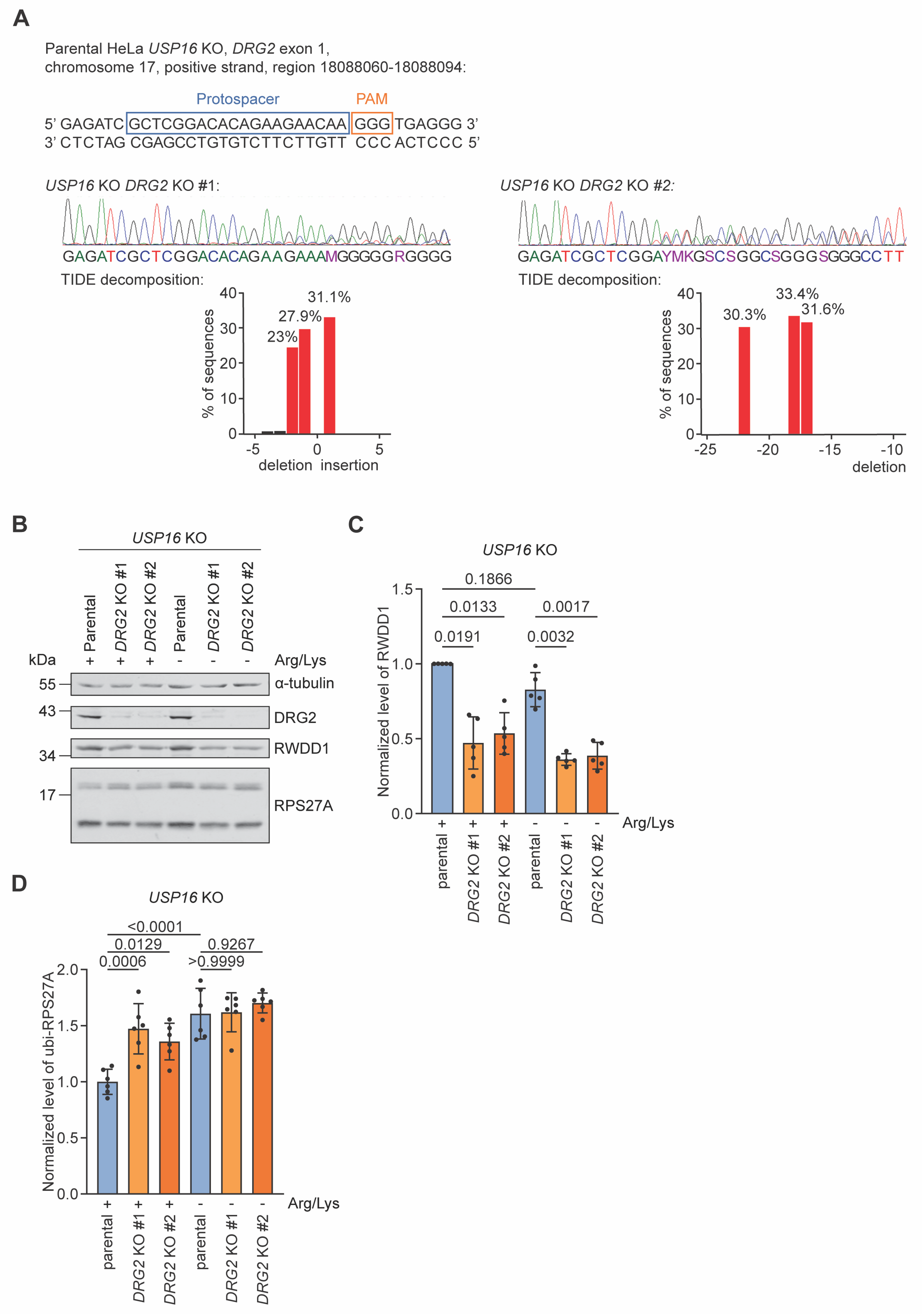


**Supplemental Figure S5. Level of ubiquitinated RPS27A/eS31 increases in the absence of DRG2.**

(**A**) Validation of HeLa *USP16/DRG2* DKO cell lines, generated using the CRISPR/Cas9 system in parental *USP16* KO cells. At the top, the target region in the exon 1 of the *DRG2* gene in parental cells is represented. Positions of protospacer and

PAM site are indicated. Below, Sanger sequencing chromatograms of the two generated clones. Sequences were decomposed using TIDE web service, and respective decomposition graphs are shown.

**(B)** Analysis of RPS27A/eS31 ubiquitination in HeLa *USP16* KO cells upon deletion of DRG2. The *DRG2* gene was deleted from *USP16* KO cells by CRISPR/Cas9. The parental *USP16* KO cells and two clones of *USP16*/DRG2 double knockout (DKO) cells were cultivated for 6 h in the presence (+) or absence (-) of arginine and lysine. Cell extracts were analysed by immunoblotting with the indicated antibodies.

**(C)** Quantification of levels of RWDD1 in cell lysates from (B), expressed as the ratio between RWDD1 and $\alpha$-tubulin, and normalized to samples from non-starved parental *USP16* KO cells (N = 5, mean ± SD, Welch ANOVA and post hoc Dunnett T3 test, p-values are indicated).

**(D)** Quantification of levels of ubiquitinated RPS27A/eS31 in cell lysates from (B), expressed as the ratio between ubi-RPS27A/eS31 and total RPS27A/eS31 (ubi-RPS27A/eS31 + unmodified RPS27A/eS31) and normalized to samples from non-starved parental *USP16* KO cells (N = 6, mean ± SD, one-way ANOVA and post hoc Tukey's test, p-values are indicated).
